# “Scentsor”: A Whole-Cell Yeast Biosensor with an Olfactory Reporter for Low-Cost and Equipment-Free Detection of Pharmaceuticals

**DOI:** 10.1101/2020.07.02.184457

**Authors:** Rachel A. Miller, Seryeong Lee, Ethan J. Fridmanski, Elsa Barron, Julia Pence, Marya Lieberman, Holly V. Goodson

## Abstract

Portable and inexpensive analytical tools are required to monitor pharmaceutical quality in technology limited settings including low- and middle-income countries (LMICs). Whole cell yeast biosensors have the potential to help meet this need. However, most of the read-outs for yeast biosensors require expensive equipment or reagents. To overcome this challenge, we have designed a yeast biosensor that produces a unique scent as a readout. This inducible scent biosensor, or “scentsor,” does not require the user to administer additional reagents for reporter development and utilizes only the user’s nose to be “read.” In this manuscript, we describe a scentsor that is responsive to the hormone estradiol (E2). The best estimate threshold (BET) for E2 detection with a panel of human volunteers (n = 49) is 39 nM E2 (15 nM when “non-smellers” are excluded). This concentration of E2 is sensitive enough to detect levels of E2 that would be found in dosage forms. This manuscript provides evidence that scent has potential for use in portable yeast biosensors as a read out, particularly for use in technology-limited environments.

---

There is a need for improved analytical tools suitable for detecting pharmaceuticals in technology-limited settings. One application for such analytical tools is to monitor the quality of pharmaceutical dosage forms, as it is estimated that at least 10% of all medical products sold in low- and middle-income countries (LMICs) are substandard and/or falsified.^1^ Equipping pharmacists and customs workers with a tool to test batches of drugs could prevent substandard or falsified pharmaceuticals from making their way to patients. HPLC is the gold standard for quantifying many pharmaceuticals.^2,3^ However, the initial cost of equipment, price of consumables, and lack of trained personnel make this technology out of reach for many areas. A device that is inexpensive, robust, user-friendly, stable, and portable would be ideal for monitoring pharmaceuticals in LMICs.

A technology that shows promise for pharmaceutical analysis in LMICs is whole cell yeast biosensors because yeast are inexpensive to maintain, have a vast genetic toolbox, and are hardy.^4,5^ However, many of these biosensors are not portable, require additional equipment to be read, or call for reagents which are not shelf-stable and/or expensive.^4,6-11^ We recently published a description of a whole-cell yeast biosensor that detects doxycycline with a fluorescent reporter which can be read with an inexpensive light box.^12^ However, it would be ideal to be able to utilize a reporting system that does not require any additional equipment or reagents.

To create an equipment-free whole cell yeast biosensor, we have developed an olfactory-based reporter. Scent has the potential to signal the presence of analytes, needing only the user’s nose to measure biosensor output. To our knowledge, scent was first used in a synthetic biology context as a biosensor reporter by the 2006 MIT IGEM team, who used it as a read-out for the physiological state of bacteria.^13^ Scent has not yet been used in published work as a read-out in whole cell biosensors, though some have described olfactory reporters for analytical purposes using mostly enzyme-based *in vitro* systems.^14-17^

These early studies, while foundational, had limitations. For example, few (if any) of these previous scent-based reporter systems were developed for field use or technology-limited settings, so cost and usability of the devices were rarely discussed. Additionally, the sample size of human volunteers used to validate these systems was generally small: in the reports listed, study sizes typically ranged from just 5 to 10 volunteers (no demographic data given), and internal review board (IRB) approvals for the use of human subjects were generally lacking. ^14-17^ These issues present concerns for the reproducibility of this data and are a roadblock for further development of these technologies for field use. Due to these issues, we endeavored to develop more rigorous standards for scent-based device testing and reporting while developing our own inexpensive and portable scent-based yeast biosensor.

After considering a variety of odorant options, we chose isoamyl acetate as the scent product for our yeast biosensor. Isoamyl acetate has a distinct banana smell which is non-toxic at levels that yeast produce and is unlikely to occur naturally in most samples being tested, including water, soil, or pharmaceuticals.^18^ Humans can detect isoamyl acetate scent at low levels: the threshold for detection of isoamyl acetate in water is 0.017 ppm.^19^ The naturally occurring enzyme, acetyl transferase I (ATF1), is primarily responsible for converting isoamyl alcohol to isoamyl acetate in yeast.^18^ The production of ATF1 in yeast has been studied extensively in the brewing industry for flavor production, and thus the pathway of isoamyl acetate synthesis has been well-characterized.^18,20,21^ For these reasons, ATF1 was selected to be overexpressed in yeast to test their ability to produce scent as readout in response to a stimulus (Figure 1C).

**Figure 1.**
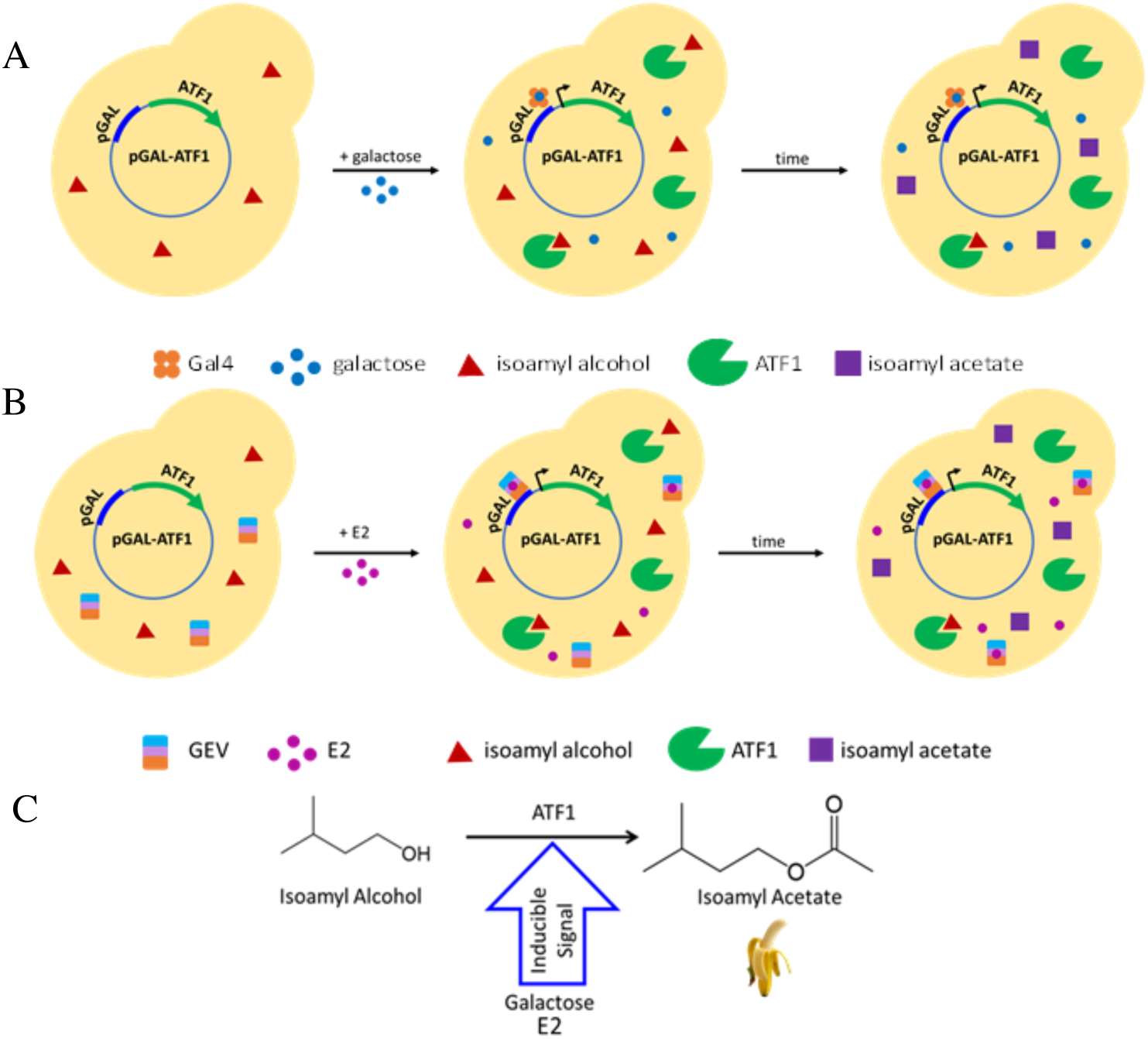
Scentsor Design. (a) Galactose scentsor made using the galactose-inducible plasmid, pATF1. When scentsor yeast are treated with galactose (in the absence of glucose), galactose binds to the endogenous Gal4 transcription factor and induces yeast to over-produce the enzyme ATF1 (note that endogenous ATF1 is present at low levels). ATF1 converts isoamyl alcohol (which yeast naturally produce) to isoamyl acetate (banana scent). (b) Estrogen scentsor made using the chimeric GEV receptor that binds to the galactose promoter and is induced by treatment with E2. (c) Schematic of the conversion of isoamyl alcohol to isoamyl acetate.

To demonstrate proof-of-principle for the suitability of ATF1 as a reporter for the presence of analyte, we chose to use the endogenous galactose-response system because the yeast galactose promoter is well-characterized and is commercially available (Figure 1A).^22,23^ Briefly, we obtained a plasmid in which the ATF1 gene was placed under control of the galactose promoter, and the galactose responsive plasmid was transformed into a strain *of S. cerevisiae* (PSY580a). Using GC-FID headspace analysis, we then measured the production of isoamyl acetate by the galactose-responsive scent-producing biosensor, or “scentsor,” in response to treatment with galactose.

In these initial experiments, we observed that galactose scentsors that were treated with galactose alone did not produce a significant amount of isoamyl acetate (Figure 2). We hypothesized that the naturally produced level of isoamyl alcohol (the isoamyl acetate precursor) in yeast was not sufficient to produce high levels of isoamyl acetate, even though ATF1 production increased. To test this hypothesis, we tried supplementing with isoamyl alcohol. After supplementing the scentsor with isoamyl alcohol, a significant amount of isoamyl acetate was produced by scentsors treated with galactose: the level of isoamyl acetate reached 115 ppm, which is 4 orders of magnitude greater than the reported human detection threshold for isoamyl acetate in water (Figure 2).^19^ Isoamyl alcohol could easily be added to packaged scentsor yeast; thus, this finding does not negatively impact the portability or usability of this device.

**Figure 2.**
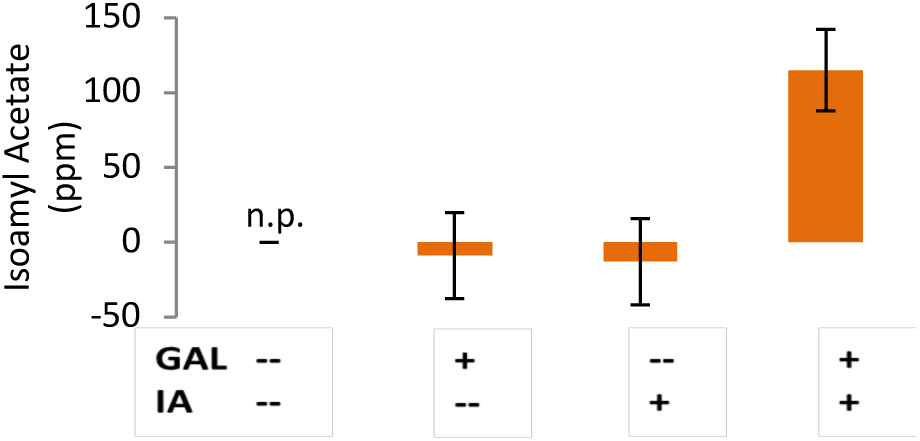
GC-FID detection of isoamyl acetate. Concentration of isoamyl acetate in the headspace of scentsor cultures was determined from a calibration curve. Error bars = Std. error from the calibration curve. Galactose scentsors were cultured in either inducing (1% galactose, 1% raffinose) or non-inducing (2% raffinose) conditions with or without added IA (250 ppm isoamyl alcohol). The abbreviation “n.p.” indicates that the Xcalibur software was unable to detect and integrate a peak for isoamyl acetate in a given run.

Because the galactose scentsor supplemented with isoamyl alcohol showed significant signal in response to galactose, we tested it with human noses (IRB: 17-12-4290).

To test the ability of humans to detect the presence vs. absence of galactose with our scentsor, we used a 3-alternative forced choice (3-AFC) model.^24,25^ Participants were presented with 5 sets of samples. Twenty-eight volunteers signed up for this study, and a total of 140 observations were collected (see Supplemental Information for more details on study size and sampling procedures). In each set of samples, there were 3 tubes of scentsors. Two of the tubes contained non-induced galactose scentsor yeast (raffinose-treated) and the other tube contained induced galactose scentsor yeast (galactose-treated). For each set, participants were asked to identify which tube smelled most like banana. Analysis of the results from this study showed that 91% of responses corresponded to the galactose-treated scentsor yeast (Table S2). Additionally, of the 28 volunteers who participated in this study, 22 of individuals chose the galactose-treated tube in all 5 of their sets (Figure 3). The Z-score for this occurrence is 63. This finding indicates that participants were able to detect the inducible banana scent produced by the yeast.

**Figure 3.**
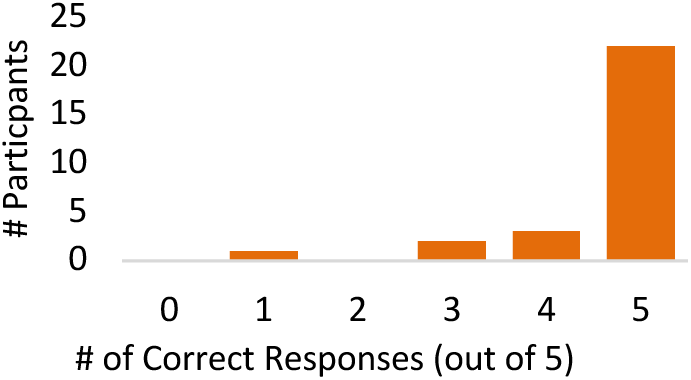
Distribution of Correct Responses from Galactose Scentsor Testing. The number of correct responses out of 5 total sets was recorded for each of the participants (n = 28).

Although most of our volunteers chose the correct tube in all 5 replicates, there were a few individuals who chose an incorrect tube in at least one of the sets, and one individual scored below the guessing probability (Figure 3). These data suggest that there are some individuals who can smell the banana reporter well, while others have weakened acuity for banana scent, if they can smell it at all. This conclusion is not surprising given that 12.4 % of the population in the U.S. was found to have a general smell impairment.^26^ Another factor that could be contributing to the failure rate is the incidence of specific anosmia (the inability to smell a particular odorant though a person’s general sense of smell is normal) for isoamyl acetate within our population. Specific anosmia can result from genetic mutations which cause the loss of receptors involved in the detection of a particular scent.^27^ We were not able to find a specific anosmia rate for isoamyl acetate in the literature, but it is known that specific anosmia can range from 0.1% to 47% for particular odorants.^28^

Because variation in performance with the scentsor was observed among panelists (as expected from the literature), a chance-corrected beta-binomial function was used to estimate the probability of true discrimination (Pd) for treated scentsors. The beta binomial is the model of choice for replicated sensory testing.^24,29-31^ This analysis shows that the probability that an individual will detect a difference in banana scent between treated and untreated samples, *P*_*c*_, is 92 ± 3% (Table S3). The true discrimination probability, *P*_*d*_, is 88 ± 5% at a 95% confidence interval(Table S3). These findings support our hypothesis that yeast biosensors could be made to produce a detectable scent in response to a given analyte, in this case galactose.

Because results with the galactose scentsor were promising, we coupled our scent reporter to a pharmaceutical detection circuit in yeast that is relevant for testing pharmaceutical quality in LMICs. We chose to develop an estrogen scentsor because sex-hormone-based drugs fall into a category of pharmaceuticals that have been found to be substandard at rates above average in LMICs: 56% of sex hormone and genitourinary drugs were found to be substandard in these regions.^32^ Additionally, there have been reported cases of substandard and/or falsified contraceptives.^33-35^

To build an estrogen scentsor, the galactose scentsor was reengineered to detect the hormone estrogen through the use of the chimeric GEV receptor.^36^ The GEV receptor binds to the galactose promoter and initiates transcription of the downstream gene upon binding to estrogen. The pGEV-TRP plasmid was transformed into the galactose scentsor to form the estrogen scentsor (Figure 1B). This estrogen scentsor was first characterized by GC-FID using the protocol established for the galactose scentsor (Figure 2). Estrogen scentsor yeast were treated with a range of estradiol (E2) concentrations and isoamyl acetate production was quantified using GC-FID (Figure 4). The estrogen scentsor had maximum production of isoamyl acetate at 1000 nM E2, reaching 125 ppm isoamyl acetate.

**Figure 4.**
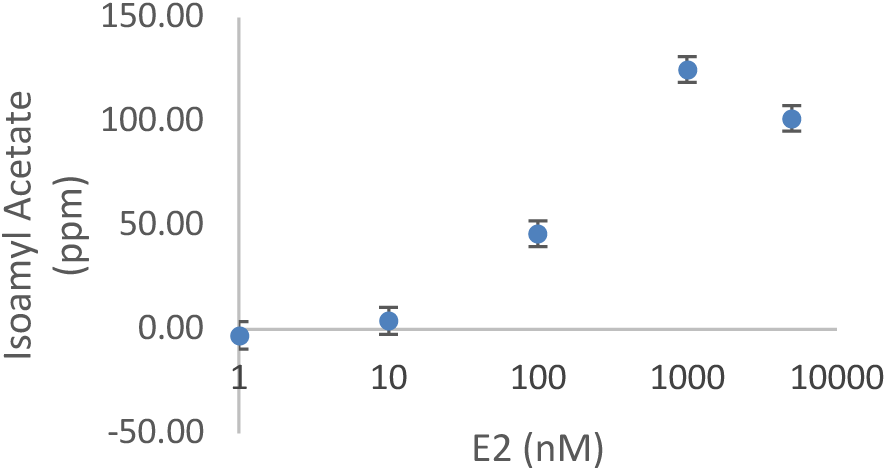
Estrogen Scentsor Dose Response. Estrogen scenstors were treated with various concentrations of E2 and allowed to incubate for 18 hrs before analysis with GC-FID. 250 ppm iso-amyl alcohol was added upon incubation. A representative experiment is shown. Error bars are the std error from the calibration curve. Note the log scale on the x-axis. 1 nM appears to drop below 0 ppm isoamyl acetate because the calibration curve does not go through the origin.

These data indicate that isoamyl acetate production rises above background with treatment of ∼10 nM E2 and reaches an EC50 above 100 nM (Figure 4). These data also show that the estrogen scentsor yeast respond to estradiol in a dose-dependent manner, except at the highest concentration of E2 (5000 nM). Isoamyl acetate production dropped in scentsors treated with 5000 nM E2, compared to those treated with 1000 nM E2. 5000 nM was chosen as the highest concentration of E2 in our experiments because it is approximately the upper solubility limit for E2 in aqueous solutions.^37^ The reason for the drop in response at high E2 concentrations is unknown, but we speculate that it could be due to an increased burden on the cells at this high level of E2. It is possible that such high concentrations of E2 either disrupt some general cell functions or the impact the production of isoamyl acetate.

Having confirmed that the GEV receptor is compatible with the ATF1 reporter gene and that estrogen scentsor yeast are responsive to estradiol (Figure 4), we determined the sensitivity of human noses for estradiol-treated scentsor yeast (IRB: 17-12-4290). The protocol for this study is based on the established methods.^25,38^ The ascending 3-AFC model was used to determine the threshold of E2 that is needed for participants to recognize the scentsor reporter signal (banana scent). Estradiol concentrations were presented in an ascending order to mitigate sensory desensitization. Participants were presented with 6 sets of estradiol-treated scentsor yeast with the following concentrations of E2: 0, 1, 10, 100, 1000, and 5000 nM E2. Each set of samples presented to participants had 2 tubes of untreated scentsor and one tube of E2 treated scentsor.

A best estimate threshold (BET) was calculated from the responses of the 49 volunteers who participated in the E2 threshold test (Table 1; Table S5). The BET is an approximation of the stimulus level detectable with a probability of 0.5 by 50% of the population. The BET for an individual participant is calculated by taking the geometric mean of the concentration at which the last miss occurred and the next higher concentration that was selected correctly. The BET for the sample population was then calculated as the geometric mean of the individual assessors.

**Table 1.** Estrogen Scentsor Results.

| E2 (nM) | 1 | 10 | 100 | 1000 | 5000 |
| --- | --- | --- | --- | --- | --- |
| Percent Correct* | 33% | 65% | 77% | 91% | 91% |

\*Excluding “non-smellers”

The BET for the detection of E2 for our panel of 49 participants was 39 nM E2 (or 10 ppm isoamyl acetate when the measured isoamyl acetate concentration was used to calculate BET) (Table S5). This concentration of E2 is sensitive enough to detect levels of E2 that would be found in dosage forms, including contraceptives, which generally contain at least 10-30 μg active ingredient (corresponding to 37-110 nM E2 if dissolved in 1 L). If non-smellers are excluded, the BET dropped to 15 nM E2 (∼ 5 ppm isoamyl acetate). Non-smellers were defined as those who got 0 or 1 answer correct out of the 5 sets that included E2 (note that the predicted number of correct answers that each participant would choose by chance is 1.6 based on our 3-AFC testing method). Given that the previously determined threshold for humans to detect isoamyl acetate in aqueous solutions is 0.017 ppm isoamyl acetate,^19^ it is likely that complex background scent of the yeast culture in which the isoamyl acetate was presented had a negative effect on their sensitivity to isoamyl acetate. However, this decrease does not significantly affect the ability of participants to detect the levels of isoamyl acetate produced by scentsor yeast when treated with concentrations of E2 that are relevant for pharmaceutical monitoring, as evidenced by the low concentration of E2 detectable by the majority of participants (Table 1; Figure 4).

Detailed demographic information is included in supplemental data (Table S4, Table S7). This information is provided so that our results may be interpreted in light of the diversity of our panel. In both studies, our panel was primarily white, female, and young when compared to other scent-based acuity studies.^26,39^ Logistic regression with these datasets suggest that the scentsor is not subject to detrimental demographic bias when in regards to usability (Table S6, Table S8). However, a larger, more diverse demographic study would need to be conducted to confirm this.

Our scentsor studies show that the probability of true discrimination with the galactose scentsor was 88%. Because there is evidence for a sub-population of non-smellers, we believe that our scentsor test could benefit from a positive control to determine whether the potential user is able to smell iso-amyl acetate. This could be presented in the form of a scratch-n-sniff sticker and would likely improve the probability of discrimination for our scentsor. This could also prevent false negative results in future studies. Additionally, the sensitivity of the GEV-based E2 scentsor does not reach the level of sensitivity needed to monitor environmental samples for EDCs. A more sensitive receptor, such as hER, would need to be used in order to reach the sensitivity needed to monitor environmental EDCs.

In conclusion, we have shown that scent producing whole cell biosensors, or scentsors, are able to produce significant amounts of isoamyl acetate upon treatment with an analyte of interest. Isoamyl acetate levels produced by scentsor yeast are detectable by the human nose. This scent-producing reporter can be coupled to estrogen sensing machinery in order to produce an estrogen scentsor. The BET for our panel of sniffers was 15 nM E2 (without non-smellers). This concentration of E2 is more than sensitive enough to detect levels of E2 that would be found in dosage forms, which generally contain at least 20 μg active ingredient. The scent-based reporter would be most useful for applications in which a threshold concentration of analyte needs to be reported, such as testing pharmaceuticals for a threshold level of API. Thus, this work shows that scent-based reporters in whole cell yeast biosensors have potential to address real-world analytical needs in technology-limited settings.

## Supporting information

Supplemental Information (materials, methods, and demographic analysis)

## ASSOCIATED CONTENT

### Supporting Information

This material is available free of charge via the Internet at http://pubs.acs.org Experimental details, yeast culturing conditions, GC calibration curve, scent study datasets, and data analysis code (PDF)

## ACKNOWLEDGMENT

Funding was provided by the Army Corps of Engineers ORISE program (RAM), the Notre Dame CBBI Training Program (JP), and the Notre Dame Department of Chemistry and Biochemistry. Some additional funding was provided by National Science Foundation DBI 1556349 (HVG). We thank the Materials Characterization Facility and the Center for Social Research at Notre Dame, which were both integral parts of this project.

## ABBREVIATIONS

LMIC: low- and middle-income countries
HPLC: high performance liquid chromatography
GC,-FID: gas chromatography-flame ionization detection
E2: estradiol
EC50: half-maximal effective concentration

## Notes

### Competing Interest Statement

The authors have declared no competing interest.

## REFERENCES

1. WHO. A study on the public health and socioeconomic impact of substandard and falsified medical products. 2017.

2. Shi YQ, Yao J, Liu F, et al. Establishment of an HPLC identification system for detection of counterfeit steroidal drugs. Journal of Pharmaceutical and Biomedical Analysis. 2008;46(4):663–669.

3. Campistron G, Coulais Y, Caillard C, Mosser J, Pontagnier H, Houin G. PHARMACOKINETICS AND BIOAVAILABILITY OF DOXYCYCLINE IN HUMANS. Arzneimittel-Forschung/Drug Research. 1986;36-2(2):1705–1707.

4. Jarque S, Bittner M, Blaha L, Hilscherova K. Yeast Biosensors for Detection of Environmental Pollutants: Current State and Limitations. Trends in Biotechnology. 2016;34(5):408–419.

5. Baronian KHR. The use of yeast and moulds as sensing elements in biosensors. Biosensors & Bioelectronics. 2004;19(9):953–962.

6. Fine T, Leskinen P, Isobe T, et al. Luminescent yeast cells entrapped in hydrogels for estrogenic endocrine disrupting chemical biodetection. Biosensors & Bioelectronics. 2006;21(12):2263–2269.

7. Leskinen P, Virta M, Karp M. One-step measurement of firefly luciferase activity in yeast. Yeast. 2003;20(13):1109–1113.

8. Weaver AA, Halweg S, Joyce M, Lieberman M, Goodson HV. Incorporating yeast biosensors into paper-based analytical tools for pharmaceutical analysis. Analytical and Bioanalytical Chemistry. 2015;407(2):615–619.

9. Jarque S, Bittner M, Hilscherova K. Freeze-drying as suitable method to achieve ready-to-use yeast biosensors for androgenic and estrogenic compounds. Chemosphere. 2016;148:204–210.

10. Chu WL, Shiizaki K, Kawanishi M, Kondo M, Yagi T. Validation of a New Yeast-Based Reporter Assay Consisting of Human Estrogen Receptors alpha/beta and Coactivator SRC-1: Application for Detection of Estrogenic Activity in Environmental Samples. Environmental Toxicology. 2009;24(5):513–521.

11. Routledge EJ, Sumpter JP. Estrogenic activity of surfactants and some of their degradation products assessed using a recombinant yeast screen. Environmental Toxicology and Chemistry. 1996;15(3):241–248.

12. Miller RA, Brown G, Barron E, Luther JL, Lieberman M, Goodson HV. Development of a paper-immobilized yeast biosensor for the detection of physiological concentrations of doxycycline in technology-limited settings. Analytical Methods. 2020;Advance Article.

13. Dixon J, Kuldell N. BIOBUILDING: USING BANANA- SCENTED BACTERIA TO TEACH SYNTHETIC BIOLOGY. Methods in Enzymology, Vol 497: Synthetic Biology, Methods for Part/Device Characterization and Chassis Engineering, Pt A. 2011;497:255–271.

14. Duncan B, Le NDB, Alexander C, et al. Sensing by Smell: Nanoparticle-Enzyme Sensors for Rapid and Sensitive Detection of Bacteria with Olfactory Output. Acs Nano. 2017;11(6):5339–5343.

15. Zhang ZY, Wang JY, Ng R, et al. An inkjet-printed bioactive paper sensor that reports ATP through odour generation. Analyst. 2014;139(19):4775–4778.

16. Xu YQ, Zhang ZY, Ali MM, et al. Turning Tryptophanase into Odor-Generating Biosensors. Angewandte Chemie-International Edition. 2014;53(10):2620–2622.

17. Mohapatra H, Phillips ST. Using Smell To Triage Samples in Point-of-Care Assays. Angewandte Chemie-International Edition. 2012;51(44):11145–11148.

18. Quilter MG, Hurley JC, Lynch FJ, Murphy MG. The production of isoamyl acetate from amyl alcohol by Saccharomyces cerevisiae. Journal of the Institute of Brewing. 2003;109(1):34–40.

19. Amoore JE, Hautala E. Odor as an Aid to Chemical Safety: Odor Thresholds Compared with Threshold Limit Values and Volatilites for 214 Industrial Chemicals in Air and Water Dilution. Journal of Applied Toxicology. 1983;3(6):272.

20. Verstrepen KJ, Van Laere SDM, Vanderhaegen BMP, et al. Expression levels of the yeast alcohol acetyltransferase genes ATF1, Lg-ATF1, and ATF2 control the formation of a broad range of volatile esters. Applied and Environmental Microbiology. 2003;69(9):5228–5237.

21. Verstrepen KJ, Derdelinckx G, Dufour JP, et al. The Saccharomyces cerevisiae alcohol acetyl transferase gene ATF1 is a target of the cAMP/PKA and FGM nutrient-signalling pathways. Fems Yeast Research. 2003;4(3):285–296.

22. Li JC, Wang S, VanDusen WJ, et al. Green fluorescent protein in Saccharomyces cerevisiae: Real-time studies of the GAL1 promoter. Biotechnology and Bioengineering. 2000;70(2):187–196.

23. Partow S, Siewers V, Bjorn S, Nielsen J, Maury J. Characterization of different promoters for designing a new expression vector in Saccharomyces cerevisiae. Yeast. 2010;27(11):955–964.

24. Bi J, Templeton-Janik L, Ennis JM, Ennis DM. Replicated difference and preference tests: how to account for inter-trial variation. Food Quality and Preference. 2000;11(4):269–273.

25. Lawless HT, Heymann H. Sensory Evaluation of Food Principles and Practices. 2 ed. New York: Springer; 2010.

26. Hoffman HJ, Rawal S, Li CM, Duffy VB. New chemosensory component in the U.S. National Health and Nutrition Examination Survey (NHANES): first-year results for measured olfactory dysfunction. Rev Endocr Metab Disord. 2016;17(2):221–240.

27. Amoore JE. Specific Anosmia: a Clue to the Olfactory Code. Nature. 1967;214(5093):1095.

28. Hirth L, Abadanian D, Goedde HW. Incidence of Specific Anosmia in Northern Germany. Human Heredity. 1986;36(1):1–5.

29. Bi J, Ennis DM. A THURSTONIAN VARIANT OF THE BETA-BINOMIAL MODEL FOR REPLICATED DIFFERENCE TESTS. Journal of Sensory Studies. 1998;13(4):461–466.

30. Bi J. Sensory discrimination tests and measurements : sensometrics in sensory evaluation. Second edition.. ed: Chichester, West Sussex : Wiley Blackwell; 2015.

31. Ennis DM, Bi J. THE BETA-BINOMIAL MODEL: ACCOUNTING FOR INTER-TRIAL VARIATION IN REPLICATED DIFFERENCE AND PREFERENCE TESTS. Journal of Sensory Studies. 1998;13(4):389–412.

32. WHO. WHO Global Surveillance and Monitoring System for substandard and falsified medical products. In: 2017.

33. WHO. Medical Product Alert No 5/2015. In. Falsified Emergency Contraceptive circulating in East Africa: World Health Organization; 2015.

34. Monge ME, Dwivedi P, Zhou MS, et al. A Tiered Analytical Approach for Investigating Poor Quality Emergency Contraceptives. Plos One. 2014;9(4):11.

35. Csillag C. Sao Paolo - Epidemic of counterfeit drugs causes concern in Brazil. Lancet. 1998;352(9127):553–553.

36. Gao CY, Pinkham JL. Tightly regulated, beta-estradiol dose-dependent expression system for yeast. Biotechniques. 2000;29(6):1226–1231.

37. Shareef A, Angove MJ, Wells JD, Johnson BB. Aqueous Solubilities of Estrone, 17β-Estradiol, 17α-Ethynylestradiol, and Bisphenol A. Journal of Chemical & Engineering Data. 2006;51(3):879–881.

38. ASTM. Standard practice E 679-19. In. Standard practice for determination of odor and taste thresholds by a forced-choice ascending concentration series method of limit. Philadelphia, PA: American Society for Testing and Materials; 2019.

39. Barber CE. Olfactory acuity as a function of age and gender: A comparison of African and American samples. International Journal of Aging & Human Development. 1997;44(4):317–334.

