## Supplemental Information (materials, methods, and demographic analysis) for "“Scentsor”: A Whole-Cell Yeast Biosensor with an Olfactory Reporter for Low-Cost and Equipment-Free Detection of Pharmaceuticals"

### Table of Contents

Materials 2

Experimental Conditions 3

Human Scentsor Studies 4

Scentsor Demographic Analysis 12

### Materials

#### *Strains and Plasmids*

S. cerevisiae strain BY4741 in which the ATF1 gene (ID:YOR377W) had been knocked-out (referred to ask "KO" in this study) was purchased from the yeast knock-out collection from Dharmacon (clone ID: 1674) (Lafayette, CO) (Table S1). The pATF1 plasmid was purchased from the Dharmacon's yeast ORF collection. This plasmid contains the GAL1 promoter upstream yeast ATF1 gene which is c-terminally tagged with Protein A, Protease 3C, HA, and His. Other plasmids and strains are indicated in Table S1.

**Table S1** List of Strains and Plasmids Used for Scent-based Biosensors

| Strain/Plasmid Name | Description | Source |
|---|---|---|
| **Yeast Strains** | | |
| PSY580a | *S. cerevisiae* Mat a; ura3Δ52, trp1Δ63, leu2Δ1, Gal2+ | Silver lab. Department of Systems Biology, Harvard Medical School (Cambridge, MA). |
| **Plasmids** | | |
| pATF1 | High copy yeast plasmid with yeast ATF1 gene under control of the GAL promoter. ATF1 gene is c-terminally tagged with Protein A, Protease 3C, HA, and His. | Dharmacon. (Lafayette, CO) |
| pGEV-Trp | Low-copy (CEN) plasmid that contains the GEV receptor under control of the constitutive MRP7 promoter with a yeast TRP selection marker. | Provided by David Eide at the department of Nutritional Sciences, University of Wisconsin-Madison (Madison, WI).[1] |

#### *Reagents and Chemicals*

Yeast nitrogen base without amino acids was purchased from BioWorld (Cat# 30626020). Amino acid dropout media was purchased from U.S. Biological (Salem, MA). Food grade isoamyl alcohol, 17-β-Estradiol (E2), and isoamyl acetate (HPLC grade), galactose, and raffinose were purchased from Sigma Aldrich. Glucose (dextrose) was purchased from Difco. GC vials (2 mL; 9 mm threads) and caps (blue with red rubber and PFTE lining) were purchased from VWR. The HP5MS column measuring 30 m x 0.25 mm x 0.25 μm film thickness was obtained from Agilent (Santa Clara, CA). SPME fibers coated with 100 μm polydimethylsiloxane (PDMS) were obtained from Supelco.

### Experimental Conditions

#### *Culturing Conditions*

To characterize the production of isoamyl acetate from the galactose-inducible plasmid (pATF1), PSY580a yeast strain was transformed with pATF1 using a standard lithium acetate method. Overnight cultures in selective medium lacking urea (SC-Ura) and containing 2% glucose were started from single colonies picked from agar plates. The next morning, overnight cultures were spun down, rinsed with sterile DI water and used to start outgrowth cultures at OD 0.6 in SC-Ura containing 2% raffinose and incubated at 30°C with shaking for approximately 6 hrs. This step was included to help deplete glucose in the yeast cultures. These cultures were then spun down and resuspended to 0.6 OD in 10 mL SC-Ura media containing 2% raffinose (non-induced) or SC-Ura 1% galactose and 1 % raffinose (induced) and placed in 50 mL centrifuge tubes. Some tubes contained 250 ppm food grade isoamyl alcohol where mentioned. Tubes were capped and incubated at 30°C with shaking for 18 hrs.

The estrogen scentsor was constructed from the PSY580a strain which was transformed with the pATF1 and pGEV-TRP plasmids. The estrogen scentsor did not require switching of metabolic carbon sources, so the following modified protocol was used when treating this GEV-based scentsor. Cultures of the estrogen scentsor were started from single colonies and grown overnight in SC-Ura-Trp media with 2% glucose. These overnight cultures were diluted to OD 0.1 in fresh selective media. The diluted cultures were treated with isoamyl alcohol and E2 to the desired final E2 concentration. Tubes were capped and incubated at 30°C with shaking for 18 hrs.

#### *Isoamyl Acetate Quantification using Gas Chromatography*

After treatment and incubation, 500 µL of scentsor culture was aliquoted into 1 mL GC vials and sealed with polytetrafluoroethylene (PTFE) lined septum caps. The samples were equilibrated at room temperature for 30 minutes before analysis. Volatiles in the headspace were extracted using a SPME fiber coated with 100 µm polydimethylsiloxane (PDMS) (Supelco, Bellefonte, PA) for 10 minutes at room temperature before being injected onto a GC-FID. Standards of isoamyl acetate and isoamyl alcohol were made dissolving each in water in 5-mL glass scintillation vials with PTFE-lined caps. Standards were stored at 4°C when used for stability studies. Isoamyl acetate standards ranged from 10-250 ppm. Standards were run on the GC and peaks were integrated using Xcalibur software (Thermo Fisher Scientific, Waltham, MA). Isoamyl acetate standards were run (n=4) and a calibration curve was constructed.

Gas chromatography-flame ionization detector (GC-FID) analysis was performed using a Thermo Trace GC coupled to an FID detector (Thermo Fisher Scientific, Trace GC ultra, Waltham, MA). The GC was equipped with a HP5MS column measuring 30 m x 0.25 mm x 0.25 µm film thickness (Agilent, Santa Clara, CA). The carrier gas used was ultra-high purity (UHP) helium with a constant flow of 2 mL/min. Oven temperature was held for 2 min at 40°C and then increased 20°C/min to 220°C and held for 5 min. Injection mode was splitless for 5 min and the injection temperature was 240°C. The flame base temperature was 240°C. The gas mixture to the flame was made up of 30 mL/min UHP nitrogen, 35 mL/min UHP hydrogen, and 350 mL/min air. The SPME fiber was injected into the inlet and volatiles allowed to desorb the length of the protocol (15 min). The fiber was cooled for 5 min before being used for the next sample. Peaks were integrated using Xcalibur software and concentrations determined from the calibration curve.

### Human Scentsor Studies

The following methods were approved by the Notre Dame Internal Review Board (Protocol ID: 17-12-4290). Volunteers were solicited through the use of e-mails, flyers, and personal interactions.

#### *Testing Procedure*

After volunteers agreed to participate in the study, they signed a consent form and were directed to the testing room. Panelists were instructed to follow the directions on a given instruction sheet. For the initial galactose-based scentsor test described below, the goal of the study was to determine if the scent response produced by treated yeast was significant from untreated yeast. A 3-alternative forced choice method was used to determine participant's discrimination ability.[2,3]

#### *Initial Galactose Scentsor Study*

Researchers presented panelists with 5 sets of 3 randomized tubes. Each set contained two vials with untreated scentsor yeast, and one tube of treated scentsor yeast. Test subjects were instructed to sniff the tubes and choose which tube contained the strongest banana scent. Participants were provided with a tube of coffee grounds and asked to cleanse their nasal palates before sniffing each set of scentsor tubes. Participants recorded their responses on a form on a computer that was provided and filled out additional demographic questions that were used in downstream analysis.

#### *Estrogen Threshold Scentsor Study*

The following scent study was developed to determine the threshold level of analyte treatment (E2 in this case) needed for participants to detect the reporter signal from blank controls. For this test, an ascending 3-alternative forced choice test was used.[4]

Panelists were instructed to follow the directions on a provided instruction sheet. Panelists were presented with 6 sets of three randomized tubes: two vials with untreated scentsors, and one tube that was the test condition. The test condition consisted of scentsor treated with E2. Sets of tubes were presented to subjects in ascending order, with the test condition with least banana scent (lowest E2 concentration) presented first and increasing concentrations of E2-treated scentsors presented as the test progressed. The final concentrations of E2 used in this study were: 0, 1, 10, 100, 1000, 5000 nM E2. Participants were instructed to sniff a provided tube of coffee grounds in order to cleanse their nasal palates before sniffing each set of scentsor tubes. Participants recorded their responses on a form on a computer that was provided and filled out additional demographic questions that were used in downstream analysis.

RESPONSES OF EACH OF THE PANELISTS WERE SCORED AS CORRECT WHEN THE E2 TREATED SCENTSOR WAS CHOSEN AND INCORRECT WHEN A NON-TREATED TUBE WAS CHOSEN. THE 0 NM "TREATED" YEAST WERE INCLUDED AS A NEGATIVE CONTROL AND THE CORRECT ANSWER DESIGNATION IS MEANINGLESS FOR THIS SUBSET OF THE DATA AND WAS THEREFORE

EXCLUDED FROM THE BET CALCULATION. THE INDIVIDUAL ESTIMATE THRESHOLD (IET) WAS CALCULATED FOR EACH PANELIST AS THE GEOMETRIC MEAN OF THE CONCENTRATION OF LAST INCORRECT RESPONSE AND THE CONCENTRATION OF THE NEXT CORRECT RESPONSE (TABLE S5

**Table** ). The BET for the panel was calculated as the geometric mean of all the IETs (Table S5). Where it was necessary to average values beyond the limits of tested concentrations (e.g. the participants did not get any successive answers correct, or they got all answers correct), the next logical step in concentration was used as a placeholder according to the method described in ASTM E679 (participants were not re-tested). The concentration used for the next logical steps in the minimum and maximum concentration of E2 (that would have been used) were 0.1 and 10,000 nM, respectively.

***Determination of Sample Size***

Previous studies that have tested scent-based sensing technologies have used very small sample sizes to validate their tests (5-10 individuals) and have thus not been able to make strong claims to the validity of their testing method. I have determined that the sample size (m) needed to estimate the proportion of people who will be able to use the technology correctly, $\theta$, within 15% accuracy ($\mu$) to be 28 at the 95% confidence interval using the following binomial error function:[5]

$$m = \frac{4\theta(1-\theta)}{\mu^2}$$

The minimum number of participants recruited for the initial galactose study was set to 30 in case some participants decide to withdraw from the study. This sample size is sufficient for estimating the preliminary usability of the scentsor. When estimating the threshold of scent a population is able to detect, a more desirable sample size is 50. ASTM method E679, which describes a method for determining the scent threshold of a population of intermediate size (50-100 participants), was used to interpret the results of the E2 threshold study.[4] This larger sample size gives a more accurate representation of the variation in the test population and allows for a more accurate estimate of the usability of the scentsor. Thus 50 was the lower limit of participants recruited for determining the threshold of analyte needed for our test population to be able to detect banana scent production.

***Results of Human Scentsor Studies***

**Table S2** Galactose Scentsor Study
x corresponds to the number of correct answers each participant made
n corresponds to the number of sets presented to each panelist

| *Judge #* | *x* | *n* |
|---|---|---|
| 4 | 5 | 5 |
| 5 | 5 | 5 |
| 3 | 5 | 5 |
| 6 | 5 | 5 |
| 7 | 5 | 5 |
| 8 | 5 | 5 |
| 9 | 5 | 5 |
| 11 | 5 | 5 |
| 10 | 5 | 5 |
| 12 | 5 | 5 |
| 13 | 5 | 5 |
| 14 | 5 | 5 |
| 15 | 5 | 5 |
| 16 | 5 | 5 |
| 17 | 5 | 5 |
| 19 | 4 | 5 |
| 18 | 5 | 5 |
| 20 | 5 | 5 |
| 22 | 4 | 5 |
| 21 | 5 | 5 |
| 23 | 3 | 5 |
| 24 | 5 | 5 |
| 28 | 5 | 5 |
| 29 | 5 | 5 |
| 30 | 4 | 5 |
| 31 | 5 | 5 |
| 32 | 1 | 5 |
| 33 | 3 | 5 |

**Table S3** Beta-Binomial Metrics for the Galactose Scentsor

| | **Definition** | **Estimate** | **Std. Error** |
|---|---|---|---|
| µ | Mean of p, probability of discrimination | 0.88 | ± 0.05 |
| γ | Measures the variation of p (over-dispersion) on the scale of probability of discrimination | 0.46 | ± 0.21 |
| Pc | Probability of a correct answer | 0.92 | ± 0.03 |
| Pd | Probability of discrimination | 0.88 | ± 0.05 |
| d’ | Distance between Signal and Signal+Noise | 2.4 | ± 0.30 |

**Table S4** Galactose Scentsor Study Results

| Demographics | n | Fraction of test population | % correct answers |
|---|---|---|---|
| **Gender** | | | |
| Female | 24 | 0.86 | 95.0 |
| Male | 4 | 0.14 | 75.0 |
| Other | 0 | 0 | N/A |
| Prefer not to say | 0 | 0 | N/A |
| **Age** | | | |
| 18-25 | 19 | 0.68 | 89.5 |
| 26-50 | 9 | 0.32 | 97.8 |
| 50+ | 0 | 0 | N/A |
| Min | 18 | N/A | |
| Max | 45 | N/A | |
| **Ethnicity** | | | |
| White | 22 | 79 | 96.4 |
| Black or African American | 1 | 0.04 | 100.0 |
| American Indian or Alaska Native | 0 | 0 | N/A |
| Asian | 4 | 0.14 | 70.0 |
| Native Hawaiian or Pacific Islander | 0 | 0 | N/A |
| Other | 1 | 0.04 | 80.0 |
| **Tobacco Use** | | | |
| Yes | 0 | 0 | N/A |
| No | 28 | 1 | 92.1 |

**Table S5** BET for Estrogen Scentsor

| | E2 (nM) | | | | | | IET* (nM E2) | IET (ppm isoamyl acetate) |
|---|---|---|---|---|---|---|---|---|
| **Judge** | **0** | **1** | **10** | **100** | **1000** | **5000** | | |
| 1 | 0 | 0 | 0 | 0 | 0 | 0 | 7071 | 145 |
| 2 | 0 | 0 | 0 | 0 | 0 | 0 | 7071 | 145 |
| 3 | 0 | 0 | 1 | 0 | 1 | 1 | 316 | 79 |
| 4 | 0 | 0 | 1 | 0 | 1 | 1 | 316 | 79 |
| 5 | 0 | 0 | 1 | 0 | 0 | 0 | 7071 | 145 |
| 6 | 0 | 0 | 1 | 0 | 1 | 0 | 7071 | 145 |
| 7 | 0 | 0 | 1 | 0 | 0 | 1 | 2236 | 116 |
| 8 | 0 | 0 | 0 | 0 | 0 | 1 | 2236 | 116 |
| 9 | 1 | 0 | 0 | 1 | 0 | 1 | 2236 | 116 |
| 11 | 1 | 1 | 0 | 0 | 0 | 1 | 2236 | 116 |
| 12 | 0 | 0 | 1 | 1 | 1 | 1 | 3 | 0 |
| 13 | 0 | 0 | 1 | 1 | 1 | 1 | 3 | 0 |
| 14 | 0 | 1 | 1 | 1 | 1 | 1 | 0.3 | 0 |
| 16 | 0 | 0 | 0 | 0 | 1 | 1 | 316 | 79 |
| 17 | 1 | 1 | 1 | 1 | 1 | 1 | 0.3 | 0 |
| 18 | 0 | 0 | 1 | 1 | 1 | 1 | 3 | 0 |
| 19 | 0 | 1 | 1 | 1 | 1 | 1 | 0.3 | 0 |
| 20 | 0 | 0 | 1 | 1 | 1 | 1 | 3 | 0 |
| 21 | 0 | 0 | 0 | 1 | 1 | 1 | 32 | 18 |
| 22 | 0 | 0 | 0 | 1 | 1 | 1 | 32 | 18 |
| 23 | 0 | 0 | 0 | 1 | 1 | 1 | 32 | 18 |
| 24 | 0 | 0 | 1 | 1 | 1 | 1 | 3 | 0 |
| 25 | 0 | 0 | 0 | 1 | 1 | 1 | 32 | 18 |
| 26 | 1 | 0 | 0 | 1 | 1 | 1 | 32 | 18 |
| 27 | 0 | 0 | 0 | 1 | 1 | 0 | 7071 | 145 |
| 28 | 1 | 0 | 0 | 0 | 1 | 1 | 316 | 79 |
| 29 | 1 | 1 | 1 | 1 | 1 | 1 | 0.3 | 0 |
| 30 | 0 | 0 | 0 | 0 | 0 | 1 | 2236 | 116 |
| 31 | 1 | 1 | 0 | 1 | 1 | 1 | 32 | 18 |
| 32 | 1 | 1 | 1 | 1 | 1 | 1 | 0.3 | 0 |
| 33 | 1 | 0 | 0 | 0 | 0 | 0 | 7071 | 145 |
| 34 | 0 | 0 | 1 | 1 | 1 | 1 | 3 | 0 |
| 35 | 1 | 0 | 1 | 1 | 1 | 1 | 3 | 0 |
| 36 | 0 | 0 | 1 | 1 | 1 | 1 | 3 | 0 |
| 37 | 1 | 1 | 1 | 1 | 1 | 1 | 0.3 | 0 |
| 38 | 0 | 0 | 1 | 1 | 1 | 0 | 7071 | 145 |
| 39 | 0 | 0 | 1 | 1 | 1 | 0 | 7071 | 145 |
| 40 | 0 | 1 | 0 | 0 | 1 | 1 | 316 | 79 |
| 41 | 0 | 0 | 1 | 1 | 1 | 1 | 3 | 0 |
| 42 | 0 | 1 | 1 | 1 | 1 | 1 | 0.3 | 0 |
| 43 | 0 | 0 | 1 | 1 | 1 | 1 | 3 | 0 |
| 44 | 1 | 0 | 1 | 1 | 1 | 1 | 3 | 0 |
| 45 | 0 | 1 | 0 | 0 | 1 | 1 | 316 | 79 |
| 46 | 0 | 0 | 1 | 1 | 1 | 1 | 3 | 0 |
| 47 | 0 | 0 | 1 | 1 | 1 | 1 | 3 | 0 |
| 48 | 1 | 1 | 1 | 1 | 1 | 1 | 0.3 | 0 |
| 49 | 1 | 1 | 0 | 1 | 1 | 1 | 32 | 18 |
| 50 | 0 | 0 | 1 | 1 | 1 | 1 | 3 | 0 |
| 51 | 0 | 1 | 0 | 0 | 0 | 1 | 2236 | 116 |
| **BET**** | | | | | | | 39 | 10 |

*Individual Estimate Threshold (IET) **Best Estimate Threshold (BET)

**Table S5** BET for Estrogen Scentsor (continued). Responses of each of the panelists are showed above, where a "0" indicates that a non-treated tube (incorrect) was chosen and "1" indicates that the E2 treated scentsor was chosen (correct). Note that judge #10 and 15 are missing from the data above due to technical issues. The IET ppm values listed were calculated from corrected values of ppm calculated from the calibration curve. The values were corrected to compensate for the fact that the curve did not go through zero, which resulted in some negative ppm values at low E2 treatment level, 1 nM: the absolute value of the calculated ppm of isoamyl acetate of background was added to each ppm value calculated from the calibration curve. IETs were calculated as the geometric mean of the last missed correct concentration and the next highest concentration. The BET is then taken as the geometric mean of the IETs. Note that IET values that were calculated to be zero were represented as zero in the BET calculation. This was done because the log of zero is undefined.

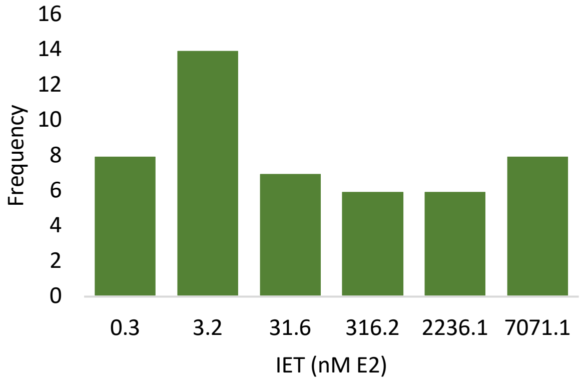


**Figure S1.** IET Histogram. The distribution of Individual Estimate Threshold (IET) frequencies for the E2 Scentsor threshold study. IETs were calculated as the geometric mean of the last missed concentration of E2 treatment and the next correct response; IETs are given in nM E2. The dataset used for this calculation is presented in Table S5.

### ***Scentsor Demographic Study***

*Method for Modeling the Impact of Demographics on Scent Test Performance: Galactose Scentsor*

We used a generalized linear model (GLM) using the "three AFC" family option from the SensR package in R in order to determine if there were significant relationships between performance in the scentsor test and collected demographic information including social constructs of ethnicity and biological differences like age, sex, and tobacco usage (see R code below). This function models logodds of a binary outcome, in our case, answering "correctly" or "incorrectly" (variable p). In this model we were interested in the probability of an outcome p for a given x value (descriptive demographic variables). The GLM models the log-odds of this outcome as a linear function of the x's using the following expression:

Logistic Regression: $\ln(p/(1-p)) = b0 + b1x1 + b2x2 + \ldots bkxk$

Where the odds ratio (OR) = $p/(1-p)$. Log odds (LO) are taken as the natural logarithm of the OR: $\ln[p/(1-p)]$. The GLM model in R returns LO values, however these are less intuitive to interpret, so the OR have been calculated. The interpretation of OR is as follows: OR > 1 = higher likelihood given a certain x value and OR < 1 = lower likelihood (p) given a certain x value. The discrete variable of scent test performance (response variable) was treated as a continuous variable in this model while all other demographic information (explanatory variables) was treated as categorical, except for age, which was also discreet. The 3-AFC family of the GLM creates a copy of the binomial family with the inverse link function changed to equal the 3-AFC psychometric function and correspondingly changed link function and derivative of the inverse link function.

*Results from Demographic Model: Galactose Scentsor*

We collected anonymous demographic data from the participants in this initial scentsor survey so that our results could be interpreted in light of the population used in our study (Table S4). Some studies have shown that women have slightly higher scent acuity in some cases and that scent acuity generally decreases with age (after 30).[6] When the 3-AFC model was run with the data that we collected, our model supported the observation that women perform slightly better than their male counterparts (Table S4; Table S6). Females have odds 3.58 greater than those of males to answer correctly which scentsor had been treated (had higher banana scent; Table S6).

Our model showed that performance increased with age over the range that was tested: for each 1 year increase in age there is a 1.15 greater odds of answering correctly (Table S6). This finding was initially surprising given that other studies saw a decrease in scent acuity with age over 30 years old.[6,7] However, the relatively young age of our panel compared to the other study could explain this difference. For this study, our panel did not contain participants over the age of 50 (the only age limitation for this study was that participants had to be older than 18 years old). Additionally, only 9 participants of the 28 fell into the 26-50 age bracket. The lack of representation in the higher age brackets could have influenced our results. A larger study would need to be conducted to determine if age impacts performance using the scentsor.

Information on ethnicity was collected to make sure the test instructions were understandable and that no implicit bias was introduced in the test procedure. Few studies have been conducted directly comparing the scent acuities of different ethnic groups. A study has linked small differences in scent acuity for particular odorants that may be linked to cultural differences and life experience.[6] Therefore, any statistical difference found between self-reported ethnic demographics could be due to bias introduced in the test procedure or due to biological differences in individuals. Our model showed that people who described themselves as Asian had 0.188 the odds of answering correctly than those who selected White (Table S6). While the p value returned from this estimate is less than 0.001, one must keep in mind the relatively small sample size used in this model when interpreting this piece of data. A larger study would need to be conducted to answer the question as to whether performance with the scentsor is subject to bias dependent reported ethnic groups.

**Table S6**. Summary of Logistic Regression of Demographic Data: Galactose Scentsor

| Model | Unit/Test Category | log Odds | Std error | Odds Ratio | Significance level |
|---|---|---|---|---|---|
| Age | Per year increase | 0.13700 | 0.06496 | 1.15 | * |
| Gender (compared to males) | Females | 1.276 | 0.473 | 3.582282 | ** |
| Ethnicity (compared to White) | Asian | 1.6706 | 0.4857 | 0.188 | *** |
| | Black or African American | 5.1082 | 560.6451 | -- (165.3724) | Not Significant |
| | Other | - 1.2562 | 0.8878 | -- (0.284734) | Not Significant |

Significance codes: p < 0 '***' 0.001 '**' 0.01 '*' 0.05 '.'

***Code used for the Galactose Scentsor Study Analysis***

The code used in R studio for the beta binomial function and the galactose scentsor GLM can be found below:

Beta Binomial Model

```
library(sensR)
library(ggplot2)
bbOutput=betabin(bb,method="threeAFC")
summary(bbOutput)
```

Logit for identifying whether demographics are predictors of performance

```
genderModel=glm(cbind(x,n-x)~relevel(gender, ref = 2) ,bb,family=threeAFC)
ageModel=glm(cbind(x,n-x)~age,bb,family=threeAFC)
raceModel=glm(cbind(x,n-x)~race,bb,family=threeAFC)
summary(genderModel)
summary(genderAgeModel)
```

***Results from Demographic Model: Estrogen Scentsor***

We performed a similar analysis of demographic data that was done for the galactose scentsor. The results from the estrogen scentsor study in regard to the demographic data collected at the time of testing was analyzed. However, because the dependent variable in this study comes from the IET values instead of count data like the galactose scentsor study, it was not straightforward to use the 3-AFC family for the GLM. A range of different E2 concentrations were presented to each panelist and so the assumption that replicates are independent of each other is violated for this sensory threshold data. Because of this, the beta-binomial model cannot be used because the calculated probability of success would not be interpretable in the conventional sense. This is why the BET calculation was used to estimate threshold concentration of stimulus needed for our panel to choose the treated samples.

However, it is reasonable to assume that participants who answered more sets of the survey correctly are likely to have higher level of acuity for banana scent. Out of 5 sets, if all of the tubes were the same (untreated) the probability of choosing a "correct" tube would be 1.67. However, we saw an elevated frequency of correct choices (corresponding to treated estrogen scentsor; Table S5). And, most of the individuals who answered more than one set correctly, did so with a pattern that suggests that correct answers were not random; correct answers were more likely to occur in "runs" at higher treatment levels than randomly dispersed across the sets (Table S5). Because of this, we believe it is possible to use the number of correct answers for a given individual to approximate the scent acuity of that individual for isoamyl acetate. In this case, a higher number of correct answers should correspond to a better scent acuity and therefore lower IET.

Assuming that the number of correct responses from each individual is representative of their IETs and therefore scent acuity for each individual, we used count data of correct responses to model the dependence of scent acuity on demographic variables. The number of correct choices for each panelist, x, was summed out of the number of sets, n. A GLM with 3-AFC family was made using this data set and the results are summarized below (

**Table** S8).

Like our previous, smaller study with the galactose scentsor which showed that females had higher odds of choosing treated scentsor yeast, females had 1.6 greater odds than males to answer correctly in the estrogen scentsor threshold study (Table S8). We plan to conduct estrogen scentsor testing in Malawi with collaborators. This study will provide a cross-cultural comparison that will help determine if this finding is universal, and thus inform test optimization. Additionally, our model showed that people who described themselves as Asian had 0.539 the odds than those who selected White (Table S8), which agrees with our galactose scentsor study. However, a larger study would need to be conducted to answer the question as to whether performance with the scentsor is subject to bias dependent reported ethnic groups.

**Table S7.** Demographic Data for Estrogen Scentsor Test

| Demographics | n | Fraction of test population | BET (nM E2) |
|---|---|---|---|
| **Gender** | | | |
| Female | 30 | 0.59 | 26 |
| Male | 18 | 0.35 | 87 |
| Other | 0 | 0 | N/A |
| Prefer not to say | 1 | 0.02 | 3.16 |
| **Age** | | | |
| 18-25 | 32 | 0.63 | 37 |
| 26-50 | 13 | 0.25 | 99 |
| 50+ | 4 | 0.08 | 3 |
| Min | 18 | N/A | -- |
| Max | 59 | N/A | -- |
| **Ethnicity** | | | |
| White | 29 | 0.57 | 22 |
| Black or African American | 1 | 0.02 | 3 |
| American Indian or Alaska Native | 0 | 0 | N/A |
| Asian | 13 | 0.25 | 247 |
| Native Hawaiian or Pacific Islander | 0 | 0 | N/A |
| Hispanic or Latino | 1 | 0.02 | 7071 |
| Other | 2 | 0.04 | 32 |
| **Tobacco Use** | | | |
| Yes | 0 | 0 | N/A |
| No | 49 | 1 | 48 |

**Table S8**. Summary of Logistic Regression of Demographic Data: Estrogen Scentsor

| Model | Unit/Test Category | log Odds | Std error | Odds Ratio | Significance level |
|---|---|---|---|---|---|
| Age | Per year increase | 0.01904 | 0.01219 | 1.0192 | Not Significant (p > 0.1) |
| Gender (compared to males) | Females | 0.4872 | 0.2211 | 1.6278 | * |
| Ethnicity (compared to White) | Asian | -0.6175 | 0.2457 | 0.539291 | * |

Significance codes: p < 0 '***' 0.001 '**' 0.01 '*' 0.05 '.'

Interestingly, there was no significant link between age and performance with the estrogen scentsor in this larger study with the estrogen scentsor (Table S8). This could be due to the fact that there were more individuals in the upper age brackets in this study compared to the previous study, but less overage of older participants as other studies which found a decrease in scent acuity with age. Another possibility is that sensitivity for isoamyl acetate is not as susceptible to age-related decline. Studies have found that acuity for different chemicals can differ across populations with very different lifestyles and so there could be biological reasons for why some odorants are more sensitive to age-related decline.[6] However, a larger study would need to be conducted to address this hypothesis.

***Code used for the Estrogen Scentsor Study Analysis***

GLM Estrogen Scentsor

```
library(sensR)
library(ggplot2)
bb_ss2=data.frame(x=rowSums(df[,2:6]),n=(rep(5,length(df$X0)))) # makes a new data frame with the
        desired variables x and n
dem$x=rowSums(df[,2:6])
dem$n=(rep(5,length(df$X0)))
genderModel=glm(cbind(x,n-x)~relevel(gender, ref = 2)  ,dem, family=threeAFC)
summary(genderModel)
ageModel=glm(cbind(x,n-x)~(Age) ,dem,family=threeAFC)
summary(ageModel)
raceModel=glm(cbind(x,n-x)~relevel(race, ref=5) ,dem,family=threeAFC) #white is reference
summary(raceModel)
```

### References


1. Gao CY, Pinkham JL. Tightly regulated, beta-estradiol dose-dependent expression system for yeast. Biotechniques. 2000;29(6):1226-1231.
2. Bi J. Sensory discrimination tests and measurements : sensometrics in sensory evaluation. Second edition.. ed: Chichester, West Sussex : Wiley Blackwell; 2015.
3. Lawless HT, Heymann H. Sensory Evaluation of Food Principles and Practices. 2 ed. New York: Springer; 2010.
4. ASTM. Standard practice E 679-19. In. Standard practice for determination of odor and taste thresholds by a forced-choice ascending concentration series method of limit. Philadelphia, PA: American Society for Testing and Materials; 2019.
5. Aitken C. Sampling - How big a sample? (vol 44, pg 750, 1999). J Forensic Sci. 2000;45(3):751-751.
6. Barber CE. Olfactory acuity as a function of age and gender: A comparison of African and American samples. International Journal of Aging & Human Development. 1997;44(4):317-334.
7. Hoffman HJ, Rawal S, Li CM, Duffy VB. New chemosensory component in the U.S. National Health and Nutrition Examination Survey (NHANES): first-year results for measured olfactory dysfunction. Rev Endocr Metab Disord. 2016;17(2):221-240.
